# An Ecological Approach to the Effects of Water-Source Locations and Time-Based Schedules on Entropy and Spatio-Temporal Behavioral Features

**DOI:** 10.1101/2021.07.15.452514

**Authors:** Varsovia Hernández, Alejandro León, Isiris Guzmán, Fryda Díaz, Martha Lorena Avendaño Garrido, Porfirio Toledo Hernández, Carlos Alberto Hernández Linares, Itzel Luna

## Abstract

In behavior analysis, the modulation of the effect of time-based schedules by the spatial characteristics of the environment has been scarcely studied. Furthermore, the spatial organization of behavior, despite its ubiquity and ecological relevance, has not been widely addressed. The purpose of the present work was to analyze the effect of water delivery location (peripheral vs. central) on the spatial organization of water-feeding behavior under time-based schedules. One group of rats was exposed to a Fixed Time 30 s-water-delivery schedule and a second group to a Variable Time 30 s schedule. For both groups, in the first phase, the water dispenser was located in the perimetral zone. In the second condition, the water dispenser was located in the central zone. Each location was presented for 20 sessions. Rat’s trajectories, distance to the dispenser, accumulated time in regions, and entropy measures were analyzed. A differential effect of the location of water delivery in interaction with the time-based schedule was observed on all the analyzed spatial qualities of behavior. The findings are discussed in relation to the ecological proposal of Timberlake’s behavioral systems.

---

The relevance of animal movement in the study of behavior was recognized in early works within the experimental analysis of behavior (Skinner, 1938). Nevertheless, the analysis of local, discrete responses with the predominant measure of response rate and the use of small experimental chambers, named as a whole *the single discrete response paradigm* (Henton & Iversen, 1978), has been the dominant methodological approach to date, with only a few exceptions (Pear, 1985; Timberlake & Lucas, 1985).

Consequently, the predominance of the single discrete response paradigm, which does not consider trajectory patterns of animals in space in the organization of behavior, has usually disregarded the spatial features of the environment. Both, the arrangement of operanda in the experimental chamber (e.g., the height of a response lever, the distance between levers) and features of the experimental arena (e.g., open vs. close zones), as spatial features related to the schedules of reinforcement (e.g. location of water or food delivery), have rarely been considered as relevant variables in the organization of animal behavior (see as exceptions the work of León, Hernández, et al., 2020 and Timberlake & Lucas, 1985; and research derived from Gibson’s ecological perspective such as Heyser & Chemero, 2012 and Jiménez et al., 2017).

By contrast, experimental psychology with an ecological approach (Timberlake, 1993; 1994) has highlighted the convenience of attending carefully to the ecological relevance of the experimental settings and observed responses to improve the parsimonious and explanatory power of the experimental analysis of behavior. Under this perspective, the temporal and spatial features of the schedule of reinforcement and the spatial setting of the apparatus could control different spatio-temporal patterns of behavior, all this within the framework of its ecological function (e.g., feeding).

From an ecological approach, animal behavior always takes place in a heterogeneous environment with an *instigation function*, given the behavioral dispositions of the animal and certain *motivational modes* (Timberlake, 1993;1994). In this sense, the spatial setting of the experimental chamber or arena is not neutral.

For example, Yaski et al. (2011) found that rats in an open field arena concentrated their trajectory patterns in the peripheral zones compared to the central zones. They also found a higher velocity in the rat’s movement in the central zone rather than in the peripheral zone. Also, rats spent more time in closed zones (i.e., peripheral zones and corners) than in open zones (i.e., central zone). For these authors, different zones of the open field arena have different ecological relevance. The closed zones serve as a refuge, safe area or home-base (Eilam & Golani, 1989; Oltmanns et al., 2021) while the central zone is an insecure area for rodents. These findings are consistent with broad literature, even in modified open-field arenas (Martinez & Morato, 2004; Alstott & Timberlake, 2009).

In addition, from an ecological approach to reinforcement (Timberlake, 1994; 2000), it has been suggested that in a situation of food or water delivery under a particular schedule to a food or water-deprived animal, a *feeding behavior system* takes place. That includes a *predatory subsystem*, *motivational modes* (i.e., general search, focal search and consume), and *perceptual-motor modules* (e.g., travel, capture, ingest, among others). From this perspective is well documented that the different motivational modes and their related perceptual-motor modules occur as a complex function of Spatio-temporal relation between stimuli and ecological dispositions related to the stimuli and the kind of conditioning procedure (Timberlake, 1983). Conversely, from the single discreate response paradigm of experimental analysis, the behavior would only be a function of the parameters and the schedule of reinforcement (Ferster & Skinner, 1957).

From a perspective in which the main variable controlling behavior is the schedule of reinforcement, delivering the same type of reinforcer with the same schedule (e.g., water delivery each 30 s), would result in the same pattern of behavior, regardless of the location of the water delivery into the experimental arena (e.g., peripheral zone vs. central zone in an open field). Contrastingly, from Timberlake’s ecological approach distinct locations of the water delivery could result in different organizations of the feeding behavior system, even under the same water-delivery schedule, if such location has a different ecological value.

In addition, from Timberlake’s ecological approach, the same stimulus (e.g., water) delivered under different schedules (e.g., water delivered with fixed or variable time) could result in a different organization of behavior, for example, related to general and focal seeking water behavior, and with its modules such as travel and its related behavioral patterns (e.g., spatial trajectories, back-and-forth patterns, variation of trajectory patterns).

Focusing on movement (or spatial behavior), studies have shown that, using time-based schedules with dispensers located in the peripheral zone of the experimental chamber, movement patterns can be modulated by the schedule of reinforcement (Eldridge et al., 1988; León, Tamayo, et al., 2020; Van Hest et al., 1986), the number of available dispensers in the experimental chamber (León, Tamayo, et al. 2020) and variations in the location of stimuli delivery. For example, in a study conducted by León, Tamayo, et al. (2020) the authors analyzed changes in movement patterns under different combinations of FT and VT schedules with water delivered in a fixed or varied location. They found more variable patterns in the combination of VT in a varied location than with FT in a fixed location.

In a more recent study, León, Hernandez, et al. (2020) conducted a study considering the previously mentioned findings. The authors analyzed spatial behavior as water delivery occurred in two different locations (in a peripheral and central zone), upon the competition of two time-based schedules (Fixed and Variable Time, Experiment 1, and 2, respectively). In Experiment 1, water was delivered according to a FT 30 s schedule. In Condition 1, water was delivered at the center of the experimental chamber; in Condition II, water was delivered close to a wall of the chamber. Each condition lasted 20 sessions, each session lasted 20 minutes.

For both schedules, FT and VT, with the dispenser located at the center of the chamber, back-and-forth patterns between peripheral and central zones were observed and the mean distance from the rat to the dispenser was higher than the distance to the dispenser when it was located close to the wall. Also, with the dispenser located at the center, the subjects concentrated more their time spent in that location and less in the peripheral zones, while with the dispenser located next to the walls the time spent by the subjects in the center was markedly lower. Also, the organisms’ variability of location was higher when the dispenser was at the center than near the wall, measured by means of an *entropy* measure. Regarding the time-based schedule, the authors found a lower distance traveled under VT with less variation of routes in comparison to the results with FT. They concluded that the locations of water delivery had differential effects on spatial behavior due to a differential ecological feature of such locations, this is, the zone close to the walls functions as a secure zone vs. the central location, considered an insecure location.

In León et al. study, the sequence of exposition to each location of the dispenser was the same for all subjects in both experiments (first at the center and then next to the wall) so it could be possible that the decrement of back-and-forth patterns and the variation of trajectory patterns and organism’s location, were due to a sequencing effect, since it is well known that the behavior variation decreases with exposure to the schedule of reinforcement through the sessions (Iversen, 2017). In order to determine if this was the case, it would be important to conduct a follow-up study in which subjects were presented with the dispenser located first next to the wall and then at the center. If the water delivery location is relevant in an ecological way, back-and-forth patterns, mean distance to the dispenser, and variability of organisms’ location (i.e., entropy) would increase with the change of water delivered from the peripheral zone to the central zone. On the other side, the opposite effect might be expected from extended exposure to the same time-based schedule if the location of water delivery has no relevance. If the location of water delivery has an effect on behavior, it would provide evidence that the spatial setting of the environment is an ecological factor that modulate behavior that is not usually considered. Regarding the effect of time-based schedules, differences in the spatial organization of water-feeding behavior, such as spatial trajectories, back-and-forth patterns, distance to the dispenser, time spent in zones, and variability of trajectory patterns and organism’s location (entropy) would be expected between fixed and variable time schedules.

The purpose of the present work is to analyze the effect of the water delivery location (wall vs. center) on the spatial organization of water-feeding behavior under time-based schedules. For a group of subjects, a Fixed Time schedule was used while for a second group a Variable Time schedule was employed.

## Method

### Subjects

Seven experimentally naïve Wistar rats were used. Rats were three and a half months old at the beginning of the experiment, each housed individually, and under a 23-hour water restriction, with free 30-min access at the end of each session. Food was freely available in their home cages. Sessions were conducted on a daily basis, seven days a week. All procedures complied with university regulations for animal use and care, and with the Mexican norm NOM-062-ZOO-1999 for Technical Specification for Production, Use and Care of Laboratory Animals.

### Apparatus

A Modified Open Field System (MOFS) was used (León, Hernandez, et al., 2020). Figure 1 shows a diagram of the apparatus. The dimensions of the chamber were 100 cm × 100 cm. All four walls of the chamber as well as the floor were made of black Plexiglas panels. Unlike the standard Open Field, in which no discreate stimuli are displayed, in the MOFS food or water can be presented in 100 different locations of the floor using the 0.8 cm holes in it, located at 0.95 cm from each other. For this study, a water dispenser, based on a servo system, made by Walden Modular Equipment® was located near the wall (Wall Condition, coordinates 95,55) or close to the center of the MOFS (Center Condition, coordinates 45,55). The central location was slightly closer to the right wall than to the left wall, this is because none of the 100 different locations for water delivery were positioned exactly at the center of the floor. When activated, it delivered 0.1 cc of water to a water cup that protruded 0.8 cm from the floor of the MOFS in one of the holes. The MOFS was illuminated by two low-intensity lights (3 watts) located above the chamber and on opposite sides of the room. Once delivered, the water remained available for 3 s for its consumption. A texturized 9 × 9 cm black patch, with 16 dots/cm, printed in a 3D printer was in proximity (5.5 cm) to the water dispenser to facilitate its location. The MOFS was cleaned using isopropyl alcohol between each experimental session.

**Figure 1.**
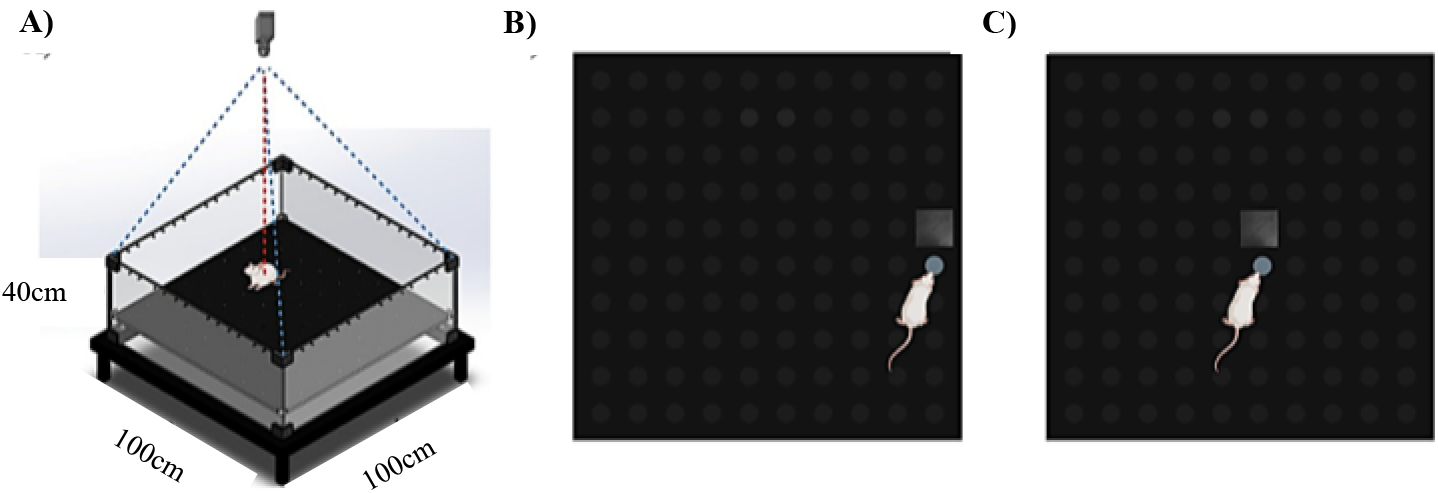
Representation of a Modified Open Field System. Note. Panel A shows an isometric view of the chamber. Panel B represents the first condition with the dispenser located next to the wall, and panel C represents the second condition with the dispenser located at the center of the chamber. The blue circle indicates the water dispenser location, and the black square represents the texturized black patch. (Created with BioRender.com)

The experimental chamber was in an isolated room on top of a 45 cm-tall table. All programmed events were scheduled and recorded using Walden 1.0 software. The rats’ movement was recorded by a Logitech C920 web camera, located at the center, 1.80 mts. above the experimental chamber. Tracking data was analyzed using Walden 1.0 software. This software recorded the rats’ location every 0.2 s in the experimental space using a system of X and Y coordinates, using their center of mass. Data files obtained from this software were then analyzed using MOTUS® and ORANGE® software.

### Procedure

Subjects were randomly assigned to one of two groups: Fixed Time (FT, 3 rats) or Variable Time (VT, 4 rats). Subjects from both groups were exposed to two consecutive conditions in the same order (see Table 1). On each condition, water was delivered using a FT 30 s schedule (Group 1) or a VT 30 s schedule (Group 2). The list of values that comprised the VT schedule were 3, 7, 13, 21, 31, 47, and 88 s; one value was randomly taken on each occasion from the list without replacement. When delivered, water remained available for 3 s. In the Wall Condition (first condition), the water dispenser was located on the floor next to a wall of the experimental chamber (see Figure 1). In the Center Condition (second condition) the water dispenser was located on the floor at the center of the experimental arena. Each condition lasted 20 sessions. Each session lasted 20 minutes.

**Table 1.** Experimental design

| Group | Time-based schedule | Condition I | Condition II |
| --- | --- | --- | --- |
| 1 | Fixed Time 30 s<br>N=3 | Wall | Center |
| 2 | Variable Time 30 s<br>N=4 | Wall | Center |

## Results

Figure 2 shows rats’ trajectories in the experimental chamber every 0.2 s for a complete session. The left panel shows data for the group under the FT schedule and the right panel for the group under the VT schedule. The first column of each panel depicts data for the last session of the Wall Condition and the second column of each panel depicts data for the last session of the Center Condition. The location of the rat in the first 0.2 s of water delivery is highlighted with a black mark. If the location of the rat at the beginning of water delivery corresponds with the location of the water dispenser, it could be an indicator that the rat was able to consume the water, nonetheless, since the water dispenser remained active for 3 s, it was still possible that rats contacted the drop of water after the first 0.2 s. Hence, the black marks correspond to the initial water capturing pattern.

**Figure 2.**
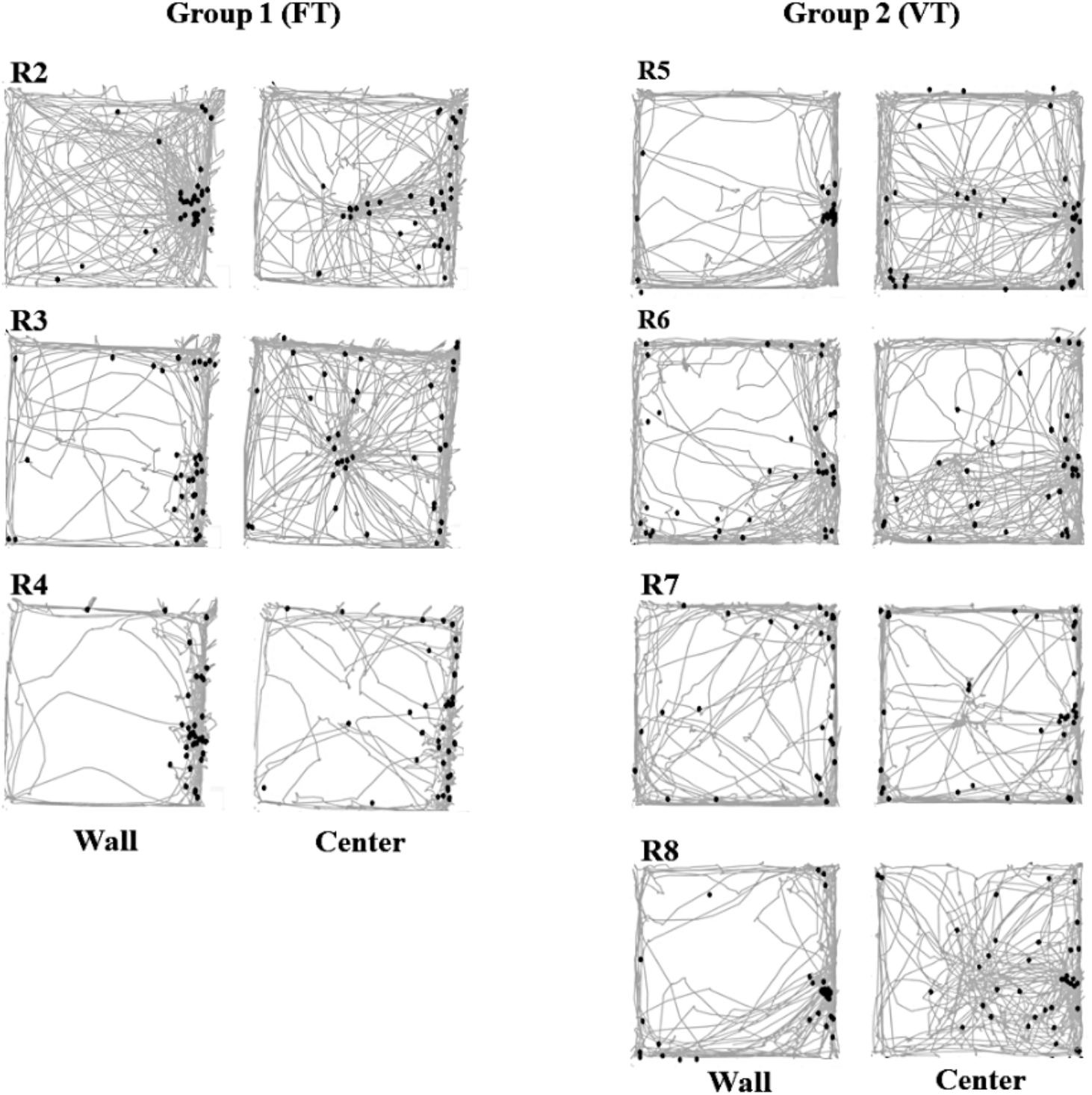
Complete routes for the last session for each rat for Condition I and II for Group 1 (left panel) and Group 2 (right panel).

For the FT group, in the Wall Condition, for all subjects, the trajectory patterns were located predominantly next to the walls of the chamber, although there were some crossings between walls. In that condition, the rats’ location at the time of delivery was close to the dispenser and predominantly on the dispenser’s wall. By contrast the Center Condition, for all rats, back-and-forth patterns between the walls, the corners, and the center of the chamber were found. For R2 and R3, their location at the time of delivery was distributed between the wall and the center of the chamber (close to the dispenser). Location of R4 at the time of delivery was away from the dispenser for most of the session. In brief, we found densification of trajectories in the dispenser zone, this is focal water-searching behavior, under both conditions.

For the VT group, in the Wall Condition, all rats showed mainly peripheral trajectories directed to the dispenser zone, although there were some crossings between walls for R6 and R7. In that condition, the rats’ location at the time of delivery was close to the dispenser located next to the wall. In the Center Condition, for all rats, distributed trajectories in the peripheral and central zone were found, directed to the dispenser, suggesting back-and-forth patterns between the walls, corners, and the dispenser zone at the center of the chamber. The location of the rats at the time of water delivery, which is the potential beginning of the water capturing pattern, was distributed between the wall and the center of the chamber (close to the dispenser).

Figure 3 shows the normalized, moment-to-moment distance every 0.2 s (grey lines) and the moving average (red lines) from the location of each rat to the dispenser. A value close to 1 means a higher distance from the rat to the dispenser and a number close to zero means minimum distance. The normalized values were obtained by assigning the value of 1 to the maximum distance value obtained for a particular session and rat and dividing each of the following values by the maximum value. The moving average was obtained using 5 time periods (for the mathematical description of this measure see Supplementary material of León, Hernández, et al., 2021). The left panel shows data for the FT group and the left panel for the VT group. Distance to the dispenser is a relevant measure because, if the location of the water dispenser (wall or center) is not relevant to the organization of water-feeding behavior, the distance to the dispenser and the back-and-forth patterns to the dispenser should remain similar among conditions. On the other hand, if the location of the water source results in different values of distance to the dispenser among conditions, this would mean that the location of the dispenser modulates the organization of water-feeding behavior.

**Figure 3.**
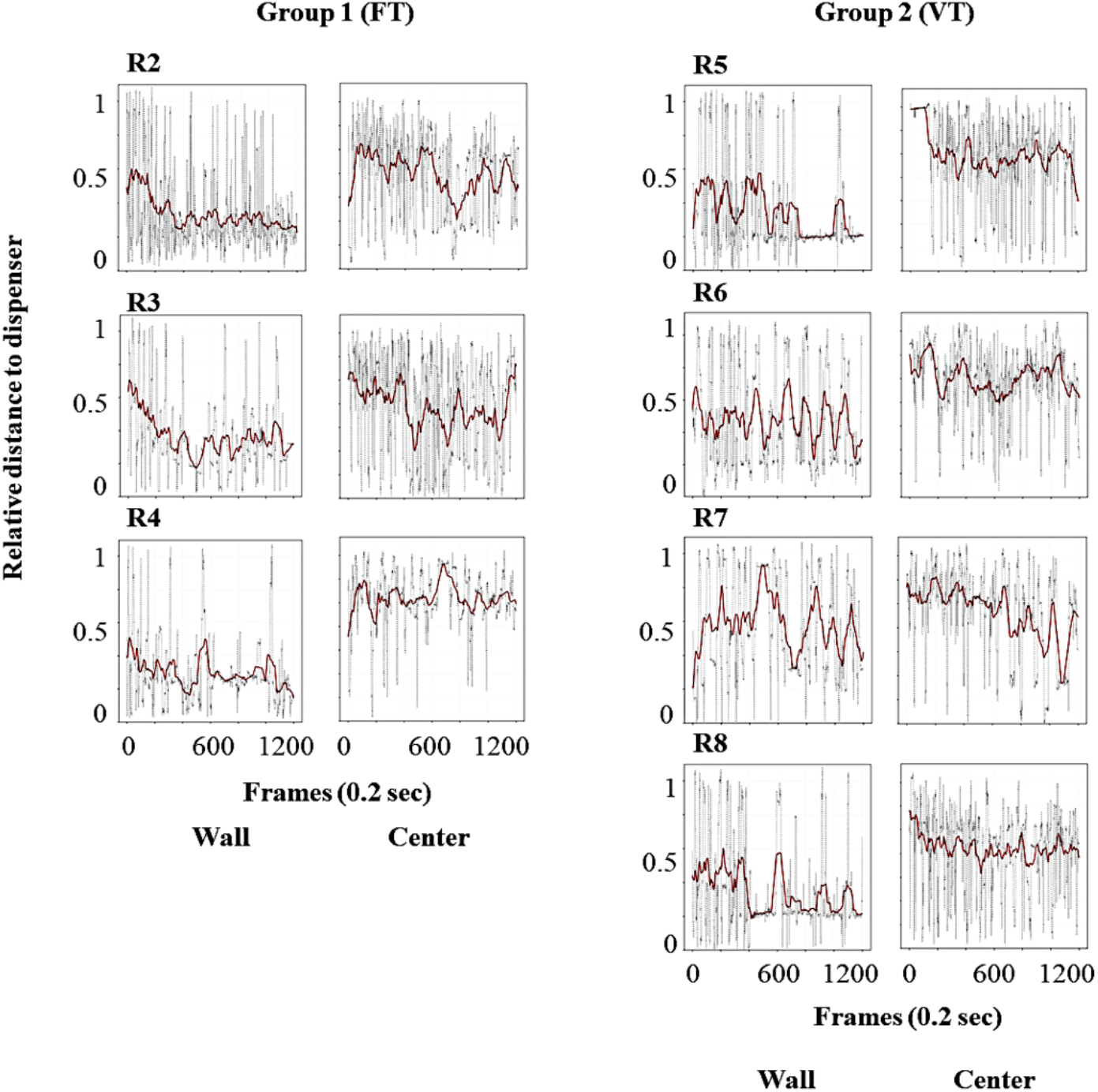
The normalized value of the distance from the rat to the dispenser every 0.2 s (gray dots) for the last session for each rat for Condition I and II for Group 1(left panel) and Group 2 (right panel).

For the FT group, in the Wall Condition, the value of the moving average of distance remained at low levels, although with some variability along the session. This result means that subjects spent most of the time close to the dispenser when it was located next to the wall. In the Center Condition, the value of the moving average remained at higher levels compared to the values obtained when the dispenser was located close to the wall, with some variability along the session; in addition, for rats R2 and R3, marked and recurring back-and-forth patterns were observed. This result shows that rats were usually far from the dispenser when the dispenser was in the center. In such a condition, rats would only approach it to drink and then move back to the walls and corners.

For the VT group, in the Wall Condition, the value of distance remained low, although with variability along the session. In the Center Condition, the distance value remained high, especially in comparison to the values obtained when it was located next to the wall. Regarding back-and-forth patterns, under the Wall Condition, they were observed only at the beginning of the session (R5, R6, and R8). By contrast, under the Center Condition, for the same subjects, well-defined back-and-forth patterns to the dispenser were observed for the whole session.

Figure 4 shows the accumulated time spent in each square region from a matrix of 10 × 10 defined virtual zones. The left panel shows data for the FT group and the right panel for the VT group.

**Figure 4.**
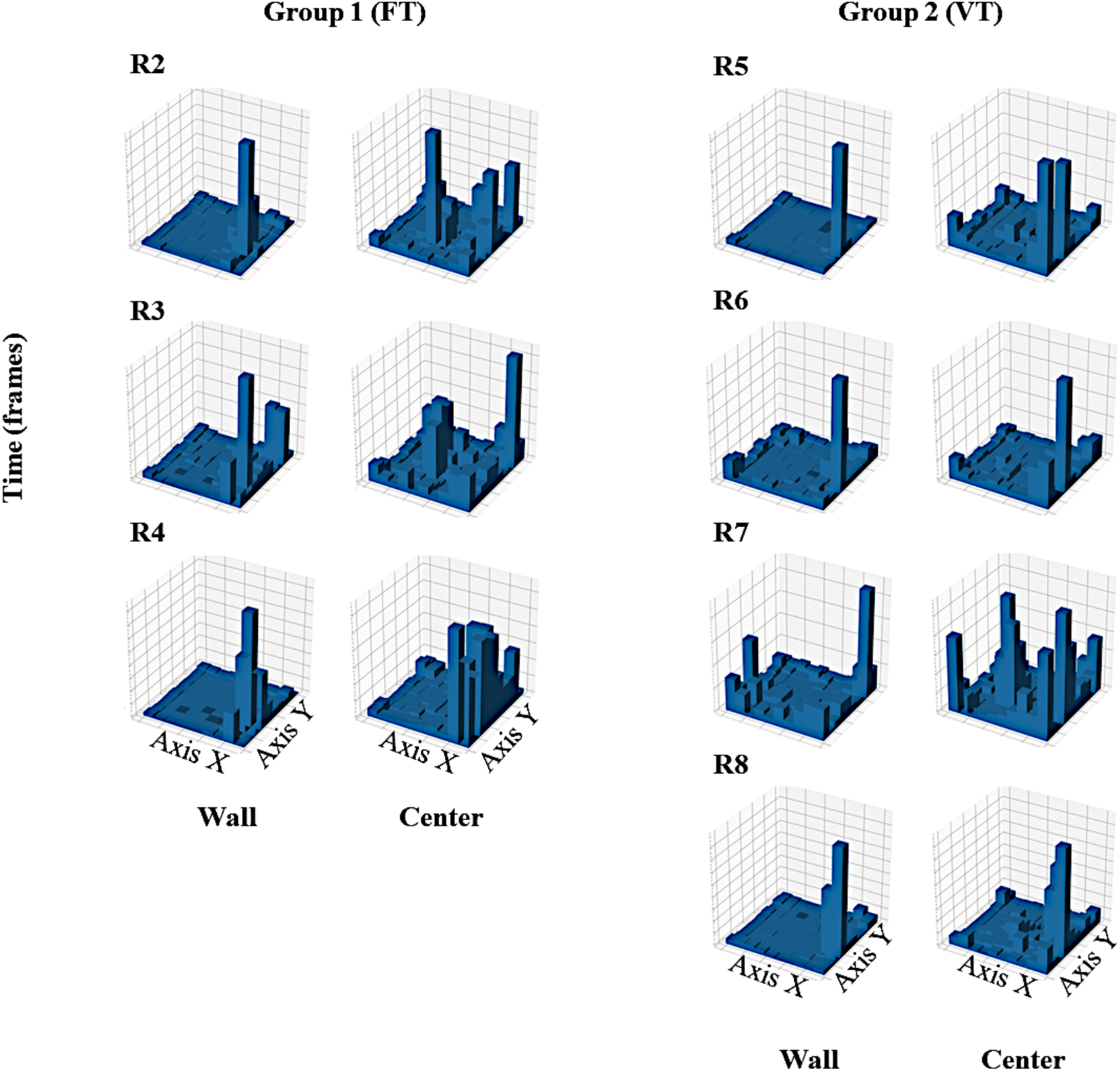
Accumulated time of stays in each of 100 zones of the MOFS for the last session for each rat for Condition I and II for Group 1 (left panel) and Group 2 (right panel).

For the FT group, in the Wall Condition, rats spent time almost exclusively close to the dispenser’s wall, mainly close to the water dispenser and to a lesser extent in the corners of the same wall. In the Center Condition, stays were distributed among the walls (mainly in the same wall of the previous phase), corners, and markedly in the center of the chamber. This result complements trajectories and distance to the dispenser analysis showing that, in the Wall Condition, rats stayed for longer periods close to the walls, while in the Center Condition the accumulated time of stays was distributed among the walls, corners, and the area of the water dispenser. A noteworthy finding is that the accumulated time of stays concentrated in the dispenser area in the Wall Condition while in the Center Condition it was spread out between the dispenser and wall areas.

For the VT group, in the Wall Condition, stays were located almost exclusively closer to the dispenser’s wall, at the center near the dispenser. Whereas the Center Condition, stays were distributed among the walls, the corners, and the center of the chamber.

The variability of location was involved in rat’s trajectories in relation to the water-seeking behavior, back-and-forth patterns to the dispenser, and the distribution of time spent in regions. To perform a quantitative comparison of the spatial dimension of behavior, as variability of location, between the two groups and conditions, we conducted an analysis of entropy of location (see Figure 5). We implemented the measure of entropy of location because it apprehends objectively and quantitatively the variability of the distribution of the spatial location of animals between the matrix of 10 × 10 defined virtual zones (see Fig 4) for a given whole session (7,200 frames).

**Figure 5.**
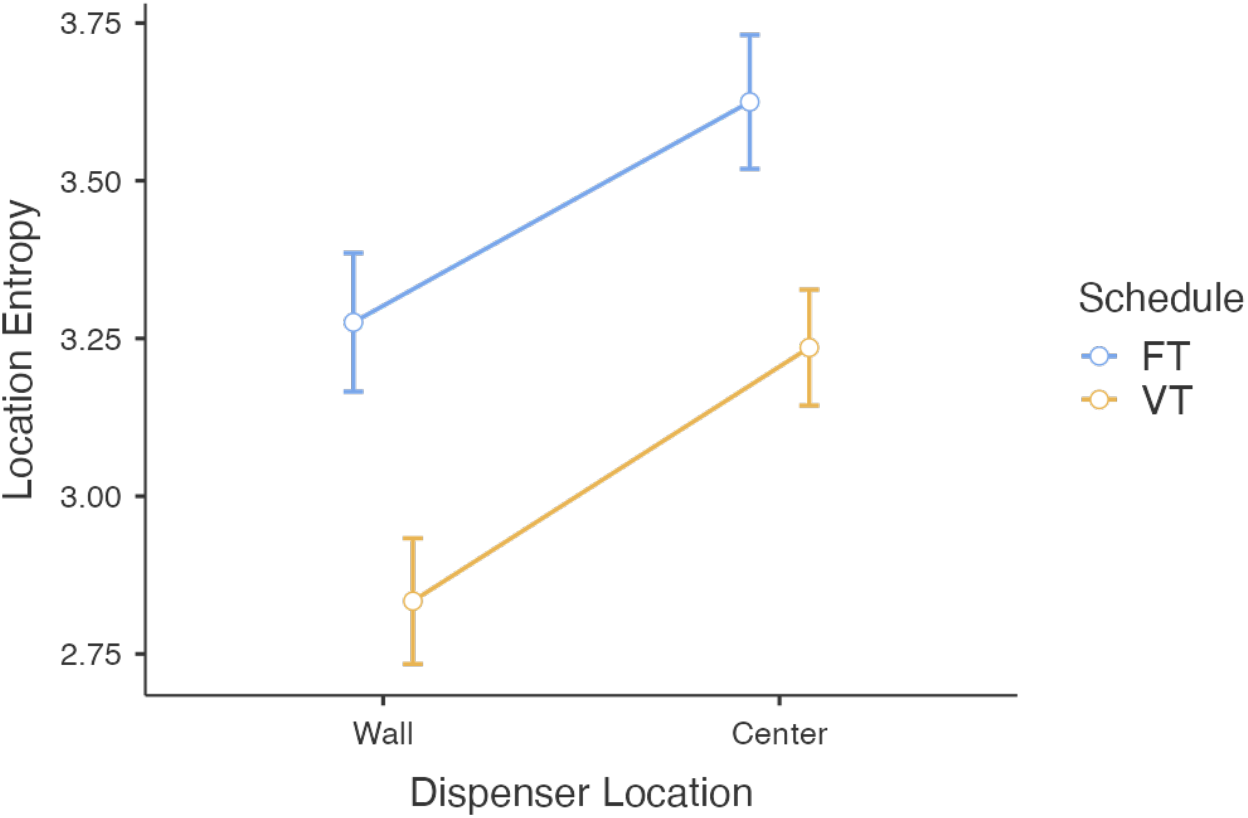
Estimated marginal means with standard error bars for location entropy for all rats in the last five sessions under each condition (wall/center) for Groups 1(FT) and 2 (VT). *Note*. Entropy for the last 5 sessions of both conditions for the three subjects for Fixed Time (blue) and Variable Time (yellow) schedules. Each plot depicts the mean (circle) and the standard error (bars), *N=66*

Shannon entropy is a measure related to a discrete random variable, which indicates variability within a distribution, it is a continuous, monotonic, and linear indicator of how different the distribution elements are from each other (Carcassi, Aidala & Barbour, 2021). Formally, given a discrete random variable *X*, in this case a given virtual region, with possible states {*x*_i_}, each with probability *P*(*x*_i_), the entropy *H*(*X*, *P*) is as follows

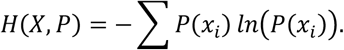

It can be proven that the entropy of a discrete random variable is a non-negative number, *H*(*X*, *P*) ≥ 0, and its measure should be maximal if all the states, this is virtual regions, are equally likely.

The discrete random variables {*x*_i_} are the permanence in each defined virtual region from a configuration of 10 × 10 defined regions, and *P*(*x*_i_) is accumulated time (standardized) spent at it.

We obtained the values of entropy of location per session for all rats in each group (3 rats for FT and 4 for VT), for the last-five sessions under each condition (5 sessions under the wall and 5 under the center dispenser location); thus, we obtained *N=70* location’s entropy data. Before conducting the analyses, we identified some outliers (*n=4*) with the Local Outlier Factor method (Breunig et al., 2000) using Orange (Demsar et al., 2013) with the following parameters: contamination: 5%, Neighbors: 20, Metric: Euclidean.

Figure 5 shows the mean value of location entropy for the last five sessions for both locations of the water dispenser and for the two groups. High values represent more variability in the organisms’ location, and low values represent less variability (León, Hernández, et al., 2020). We found higher values of entropy under the center condition than the wall condition for both schedules (FT and VT). This means that the organisms’ location was more varied with the dispenser at the center than under the wall condition. Related to reinforcement schedule, the entropy value was higher for the Fixed Time than the Variable Time schedule under both dispenser locations.

Finally, a factorial ANOVA was conducted to compare the main effects of the dispenser location condition (wall, center) and reinforcement schedule (Fixed Time, Variable Time) on entropy location. The main effects were statistically significant at the 0. 001 significance level. The main effect for dispenser location yielded an F(1, 62) = 13.5391, p < .001, indicating a significant difference on the values of entropy of location between the wall dispenser condition (M = 3.03, SD = .0818) and the center dispenser condition(M = 3.40, SD = .0770). The main effect for reinforcement schedule yielded an F(1, 62) = 16.5784, p < .001, indicating a significant difference in values of entropy of location between Fixed Time schedule (M = 3.46, SD = .0832) and Variable Time schedule (M = 3.05, SD = .0736). The interaction effect was non-significant, F(2, 62) = .0661, p > .05

## Discussion

The purpose of the present work was to analyze the effect of water delivery location (wall vs. center) on the spatial organization of water-feeding behavior under time-based schedules.

Similar to León, Hernández, et al. (2020), we found, under both schedules (FT and VT) and both conditions of water delivery location (Wall and Center), wide and diverse travel trajectories for all subjects. These trajectories could be seen as the spatial dimension of a *general search mode* associated with *travel* and *investigative* patterns in water-deprived rats. Also, we found significant densification of trajectories directed to the water dispenser, suggesting that a *focal water search* emerged for both water delivery locations under both schedules. Nevertheless, a remarkable finding is that when the dispenser was at the center zone, at the moment of water delivery, which could be seen as the beginning of the pattern of *capturing* and *consuming* water, rats were usually far and scattered from the dispenser (they were in the peripheral zone, near to the corners and walls), while when the dispenser was in the peripheral zone, the animals were near to the dispenser. This finding suggests that a *chase pattern* (i.e., approximation from a distant location to the water source once water is available) in the focal search emerged under the Center Condition, while under the Peripheral condition, a *lie-in-wait pattern* took place (i.e., wait at the location of water delivered until it is available). That is a noticeable finding because the same rats under the same schedule of water delivery showed different focal search patterns only due to the different locations of the water dispenser. This argument gains strength with the marked back-and-forth patterns to the dispensers observed under the Center Condition compared to the Peripheral condition. These findings show that the different patterns of focal search are supported by the environment’s spatial organization (Timberlake, 1993).

The distribution of time spent in regions showed that in the Wall Condition, under both schedules, rats spent most of their time in the peripheral zone, specifically in the dispenser’s wall between the dispenser location and corners. This preference for close zones is widely reported in the literature (Alstott & Timberlake, 2009; Martínez & Morato, 2004; Yaski et al., 2011; Zadicario, et al. 2005) and was intensified by the wall location of the dispenser. By contrast, under the Center Condition, both the peripheral zone (walls and corners) and the center zone accumulated high values of time spent. The observed preference for the open zone is atypical in healthy rats, even under water-deprivation conditions (León, Tamayo, et al. 2020). Thus, with the dispenser at the Center Condition, two ecological relevant environmental segments, these are feeding zone and safety zone, with associated approach patterns took place in different physical areas: the zone of water delivery at the center and the wall as the secure zone and home base (Eilam, & Golani, 1989). In contrast, under the dispenser at the Wall Condition, both functional segments (zone of water delivery and safety area) converged in the same geometrical segment: the perimetral zone.

The findings related to time spent in open vs. close zones, seen from an ecological perspective, allow understanding the emergence of the different patterns of focal water-search previously described: *chase* vs. *lie-in*-*wait* patterns. Because the open vs. close zones have different instigation functions (i.e., they are associated with different patterns), in a non-secure area is unlikely that the rat presented *stay* and *lie-in-wait* patterns of behavior, but not in a safe, close zone. So, if the water is delivered in an open zone, long excursions would take place from the secure zone (walls or corners) to *chase* the water, giving rise to back-and-forth patterns.

Based on the previous findings is evident that the different locations of the water delivery resulted, under both schedules, in distinct organizations of the water-feeding system, specifically related to focal water-searching behavior, even under the same time-based schedule. It is important to note that the findings discussed here would hardly be expected or explained from the standard single discrete response approach, but they are easily described under the ecological approach (Timberlake, 1993).

Regarding the comparison in the organization of the water-feeding behavior between Fixed Time vs. Variable Time schedules, it is challenging to identify, without deeper analysis, objective and robust differences based on trajectories, distance to the dispenser, and time spent in regions plots. Nevertheless, the entropy measure apprehends an embedded feature of the spatial behavior involved in the trajectories, back-and-forth patterns, and the time spent in regions. That is the variability of the distribution of the organism’s location. The entropy measure showed a significant difference between the Wall and Center Conditions in both schedules, strengthening the findings regarding trajectories, distance to the dispenser, and time spent in regions. But, more importantly, the entropy measure made it possible to compare, in an objective and quantitative manner, the spatial organization of water-feeding behavior under both time-based schedules. Using this measure together with analysis at the individual level (trajectories and time of stays) we could see that under the Fixed Time Schedule there was more variability in the distribution of the organism’s location than under the Variable Time Schedule. This last finding could imply differences in *general* and *focal water-searching modes* for the same dispenser location but under different time-based schedules.

The decrement of variability of rat’s location, with the dispenser located in the perimeter, was more salient under the VT schedule than under FT, an expected finding given the literature (Eldridge et al., 1988; Van Hest et al., 1986). The decrement of variability of rats’ location, with the dispenser located in the perimeter, was more salient under the VT schedule than under FT, also an expected finding given in the literature (Eldridge et al., 1988; Van Hest et al., 1986). It is plausible to suppose that this decrement in entropy of location could be associated with an increment in the *lie-in-wait* pattern in focal search given that water delivery under the Variable Time schedule is less predictable. This argument is supported by previous findings that showed that unpredicted food delivery produces a mode of *focal search* and *waiting* (Timberlake, 1986).

On the other hand, the highest entropy value was observed under FT, especially with the dispenser at the center of the arena. It has been discussed that under Fixed Time schedules, components of species-typical feeding patterns are elicited, including general search patterns (Timberlake & Lucas, 1985), these patterns imply extended trajectories (Pear, 1985) that are associated with high entropy compared to VT. The high entropy associated with FT was intensified with the water delivery at the center because it interacted with other species-typical patterns, given the structural elements in the open field, this is, the locomotion near the walls or security seeking and the avoidance of the open zones, even more so with the open field lighted (Alstott & Timberlake, 2008) as in this work. The interaction between searching, both general and focal directed at the center, with *security-seeking* patterns gave rise to the back-and-forth patterns between the water-delivery zone and secure zones as well as extended locomotion around the open field close to the walls. All these complexities and high dynamics were associated with the highest entropy.

It is worth noting that entropy values under FT in the Wall Condition and VT in the Center Condition were relatively close, this finding would be unexpected if only the time-based schedule was considered, as would be suggested from the standard approach. It is convenient to mention that although the entropy value is sensible to spatial features embedded in the water-feeding behavior system, it captures only a partial way of its temporal dimension, for this reason, it is important to supplement the entropy analysis with analyses of the Spatial-temporal organization of behavior such as the ones conducted in this work (i.e., moment to moment distance to the dispenser; rats’ location to the moment of water delivery). Future work must advance in the development and implementation of quantitative analysis that apprehends in a more comprehensive way that temporal dimension of the spatial organization of the feeding behavior system.

The present work provides robust evidence that the decrement of back-and-forth patterns to the dispenser, the concentration of stays close to the dispenser, and the decrement of entropy, were due to locating the water dispenser close to the wall, which could instigate species-typical patterns. Also, the present work shows that the decrement of back-and-forth patterns was not only an effect of an extended exposition to the same schedule of water delivery. In this sense, is noticeable that the spatial dynamics increased (i.e., back-and-forth patterns, extended trajectories, and highest entropy) in the last phase for both groups when the dispenser was at the center. As mentioned above, this is explainable by the organization of different water-search patterns due to the experimental conditions arranged.

The results of the present work add to previous findings, as no counterbalancing was considered in a previous study (León, Hernández, et al., 2020). In the present study, we conducted quantitative analyses of entropy, not carried out in previous studies, that showed a significant difference between the Wall Condition vs. Center Condition and for both groups, FT and VT schedules. On the other hand, even though Leon, Hernández, et al. (2020) expressed in the rationale of their study an ecological approach, their findings are discussed more in terms of the spatial features of the behavior than from an ecological-behavioral approach. So, the present work advance from the previous one in the following ways:

1. It strengthens the previous results under a new arrangement of conditions that allows, together with the previous findings, to control the effect of the exposure sequence and to strengthen the effect of the location of the dispenser (wall vs. center) on the organization of water-feeding behavior.
2. The implementation of quantitative analysis to identify differences between conditions and experiments that are difficult to identify with just a visual analysis.
3. The integration of a comprehensive ecological approach (Alstott & Timerlake, 2009; Timberlake, 1986, 1993, 1994; Timerlake & Lucas, 1985) to describe and discuss the spatial organization of the behavior under a water-feeding system.

The present works contribute with a novel methodological approach, based on the continuous, moment to moment, record of the center of mass in the analysis of spatial features embedded in animal movement. This approach accounts for the organization of water-feeding behavior systems such as exploration, general and focal search, among others. This approach differs from the classical ethological approach centered on ethograms that in its classical way are not automated, are discrete, and susceptible to anthropomorphic bias.

The developed approach could be strengthened with automated multi-point tracking systems for automated pose records (Datta, 2019; Mathis & Mathis, 2020), sensible measures for Spatial-temporal organization, and integrated and multidimensional analysis based on computational algorithms to characterize in a more comprehensive way the continuum of Spatial-temporal organization of water-feeding behavior system (León et al. 2021; León, 2022).

To conclude, there is pending empirical work for future studies. The current study invites us to consider spatial features of the experimental settings and their interaction with schedules of stimuli presentation (food or water) on the organization of the water-feeding system and its transitions between the motivational modes of general and focal search and handling-consumption.

As a corollary, considering that the spatial-displacement distribution is an embedded feature of different relevant behavioral phenomena such as exploration, space learning, and conditioned place preference, among others; we argue that location entropy is a promising measure to systematize findings concerning the distribution of spatial displacement of animals on surfaces. The measure of location’s entropy could be advantageous to develop an objective approach to analyzing such spatial distribution under the influence of different variables. Based on the data of the present study, we propose that location entropy could improve not only our understanding of behavior, but the reproducibility, comparability, and robustness of behavioral research based on spatial features. Therefore, future work should be aimed at evaluating this proposal.

